# ANDA: An open-source tool for automated image analysis of neuronal differentiation

**DOI:** 10.1101/2023.04.27.538564

**Authors:** Hallvard Austin Wæhler, Nils-Anders Labba, Ragnhild Elisabeth Paulsen, Geir Kjetil Sandve, Ragnhild Eskeland

## Abstract

**Background:** Imaging of *in vitro* neuronal differentiation and measurements of cell morphologies has led to novel insights into neuronal development. Live-cell imaging techniques and large datasets of images has increased the demand for automated pipelines for quantitative analysis of neuronal morphological metrics.

**Results:** We present ANDA, an analysis workflow for quantification of various aspects of neuronal morphology from high-throughput live-cell imaging screens. This tool automates the analysis of neuronal cell numbers, neurite lengths and neurite attachment points. We used rat, chicken and human *in vitro* models for neuronal differentiation and have demonstrated the accuracy, versatility, and efficiency of the tool.

**Conclusions:** ANDA is an open-source tool that is easy to use and capable of automated processing from time-course measurements of neuronal cells. The strength of this pipeline is the capability to analyse high-throughput imaging screens.

## Background

One of the defining characteristics of the central nervous system is the neuronal interconnectivity which facilitates the cell-to-cell communication required for normal brain function [1]. Establishment of neuronal networks in the developing brain are constituted by neuronal connections and can be influenced by the surrounding glia [2]. Neural development is a spatiotemporally fine-tuned biological process that spans the genesis of neurons to the maturation of functional neural tissues. The differentiation of neural cells is composed of steps such as cellular proliferation, neurite extension, neurite branching, synaptogenesis, and refinement of connections [3]. Modelling neuronal differentiation in a dish can provide new insights into how these connections are formed and altered. Many in vitro neuronal models are in use for genetic and pharmacologic screens such as neuronal differentiation cultures of mouse and human embryonic stem cells, induced pluripotent cells, neuronal stem cells, immortalized tumour cells (human neuroblastoma cells SH-SY5Y), NT2 human embryonal carcinoma cells, PC12 rat pheochromocytoma cells, and chicken and rodent primary neuronal cultures [4–15]. When microscopy is applied in these studies, the focus has been on changes to different morphologic parameters of neuronal cells, often in a high-throughput manner [16–23]. The use of label-free time-course phase contrast microscopy has increased our understanding on the rise, development, and maturation of neuronal networks without being confounded by factors such as phototoxicity [23–25]. Moreover, live-cell imaging with high spatial and temporal resolution have resulted in a massive increase in data volume and complexity [26, 27]. Image processing of large datasets from high throughput imaging platforms generally involve many steps of image pre-processing, segmentation, phenotype quantification and subsequent analysis [28, 29]. This stresses the need for software applications for large image data sets and automated approaches for reproducibility.

We have developed ANDA, a tool for automated high-throughput image analysis of live neuronal cell cultures. ANDA is a desktop application built with TAURI and features a custom made pipeline which performs image analysis in Fiji [30, 31]. We show that ANDA can quantify various metrics of three neuronal cell models with distinct differences in morphologies: chicken cerebellar granule neurons (CGNs), neuronally differentiating rat PC12 cells (PC12Ns) and pre-terminally differentiated human-derived NTERA2 cells (NT2Ns). Decisive metrics in neuronal morphology, particularly cell bodies, neurites and neurite attachment points are retrieved and reproducibly quantified, either at single time points or in time series. ANDA is open source under the MIT license and is available on GitHub (https://github.com/EskelandLab/ANDA).

## Implementation

### Image analysis of neuronal cell types

ANDA is a tool that can measure quantity, size and shape of cell bodies, neurites, and neurite attachment points from segmented images. Furthermore, ANDA fully automates image analysis and output data summarization (Figure 1). Prior to quantification, ANDA can be used to threshold raw images and apply different algorithms from Fiji [30] for segmentation. The pipeline also includes the option of using pre-segmented images as input. After thresholding and watershed algorithms are implemented, ANDA identifies cell bodies and neurites based on segmented images. This measurement requires customization of the size and circularity of the cell bodies and neurites for the neuronal cell type of interest (Supplementary Table 1). Sobel edge detection is applied to identify the neurite attachment points and all output data is summarised in csv-files.

**Figure 1.**
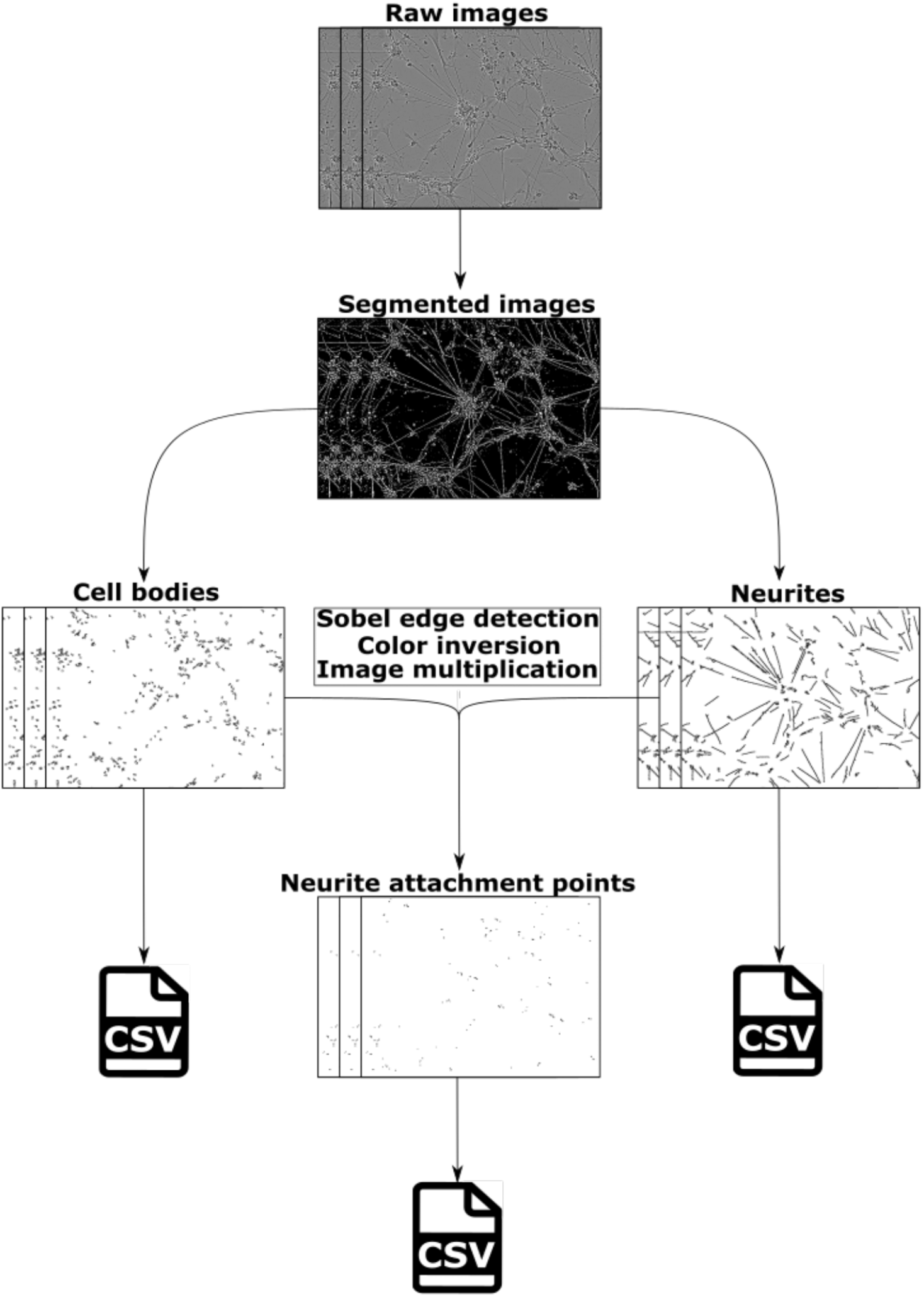
A flowchart of ANDA. Raw phase contrast images are segmented prior to identification and analysis of cell bodies or neurites. The mapping of cell bodies and neurites yield results of their own, or their outlines can be used to identify neurite attachment points. This is done by applying Sobel edge detection on the neurite outlines, followed by colour inversion of cell body outlines and neurite edges. Thereafter, the overlap between cell bodies and neurites are determined by image multiplication. These overlaps are quantified as neurite attachment points. The data is summarized in different csv files.

### Workflow

ANDA’s workflow is designed to be straightforward and easy to customize for the user. The downstream image analysis parameters are set using a graphical user interface. Before automated image analysis, settings for cell line and neurite aspect ratio inclusion threshold have to be defined as input parameters. The images of CGNs, NT2Ns, and PC12Ns are successively processed and analysed in Fiji [30] and the raw output saved in csv-files. After completion of the image analysis, mean values of each selected analysis metric for each image are summarized into separate csv-files.

### Quantification of neuronal morphological metrics

ANDA presents the option to threshold images from multiple global thresholding algorithms available in Fiji [30]. In addition, the user can choose to apply a watershed algorithm to segment the images even further or use pre-segmented images as input and thereby skip a redundant segmentation step altogether. Some cell types such as the pre-terminally differentiated NT2Ns exhibit contrast-levels that are too low to be reliably distinguished from background using standard thresholding methods, necessitating a separate segmentation-step of Weka segmentation. Weka segmentation is an unsupervised trainable machine learning algorithm that is included in Fiji, and which can improve the delineation of low-contrast objects from background, given proper training (Supplementary Figure 1)[32]. Following segmentation, the quantity, size and shape of cell bodies, neurites and neurite attachment points, are measured using built-in features in Fiji [30]. Cell bodies are isolated from background by applying image thresholding followed by a watershed algorithm. Similarly, neurites can be retrieved by isolating the motifs from background with thresholding and watershed algorithm. Cell bodies and neurites are quantified by identifying motifs with custom-set size and circularity criteria specified for each cell type. Neurite attachment points, a collective term for neurite trunks and neurite terminal ends, are retrieved by highlighting the edges of the neurite outlines with Sobel edge detector after thresholding and watershed algorithm, and thereafter isolating the overlap between cell body outlines and neurite outlines by colour inversion and image multiplication.

## Results

### Qualitative measurement assessment

Automated quantification of neuronal metrics from high-throughput experiments is the main purpose of our pipeline (Figure 1). The quality of ANDA’s ability to quantify neuronal differentiation metrics was assessed using outlines of identified structures in freshly plated and *in vitro* day 3 CGN cells (Figure 2), as well as freshly plated and differentiated PC12N (Supplementary Figure 2) and NT2N (Supplementary Figure 3). Details of the cultivation of these three cell types are described in supplementary information[18, 33]. To obtain phase contrast images of the cells we used the live-cell imaging platforms IncuCyte® ZOOM for the CGN and PC12N cells, and IncuCyte® S3 for the NT2N cells (Supplementary methods). Generally, freshly plated cells are spherical and do not exhibit neurite outgrowth until proper attachment to the growth vessel. ANDA consistently identified cell bodies in freshly plated chicken cells and *in vitro* day 3 (Figure 2B and 2F). This trend was also observed in the PC12N and NT2N models (Supplementary Figure 2B, 2E, 3B and 3E), although the NT2N cell bodies tend to have a more oblong shape. Furthermore, ANDA identified CGN neurite structures at *in vitro* day 3 (Figure 2G), with some artefactual identification of neurites in freshly plated cells (Figure 2C). ANDA detected neurite structures in three days differentiated PC12N cells but not in freshly plated cells (Supplementary Figure 2C and 2F). Some neurite structures were detected in the freshly plated NT2N cells with a clear increase after three days of differentiation (Supplementary Figure 3C and 3F). The identification of neurite attachment points relies on identified neurite and cell body structures in the image. In freshly plated CGN cells, ANDA falsely detect some neurites that also result in detection of false positive neurite attachment points (Figure 2D). True positive identification of neurite attachment points was more consistent at in vitro day 3 (Figure 2H). Based on these observations, we show that ANDA’s ability to quantify neuronal differentiation metrics improves with neuronal morphological development.

**Figure 2.**
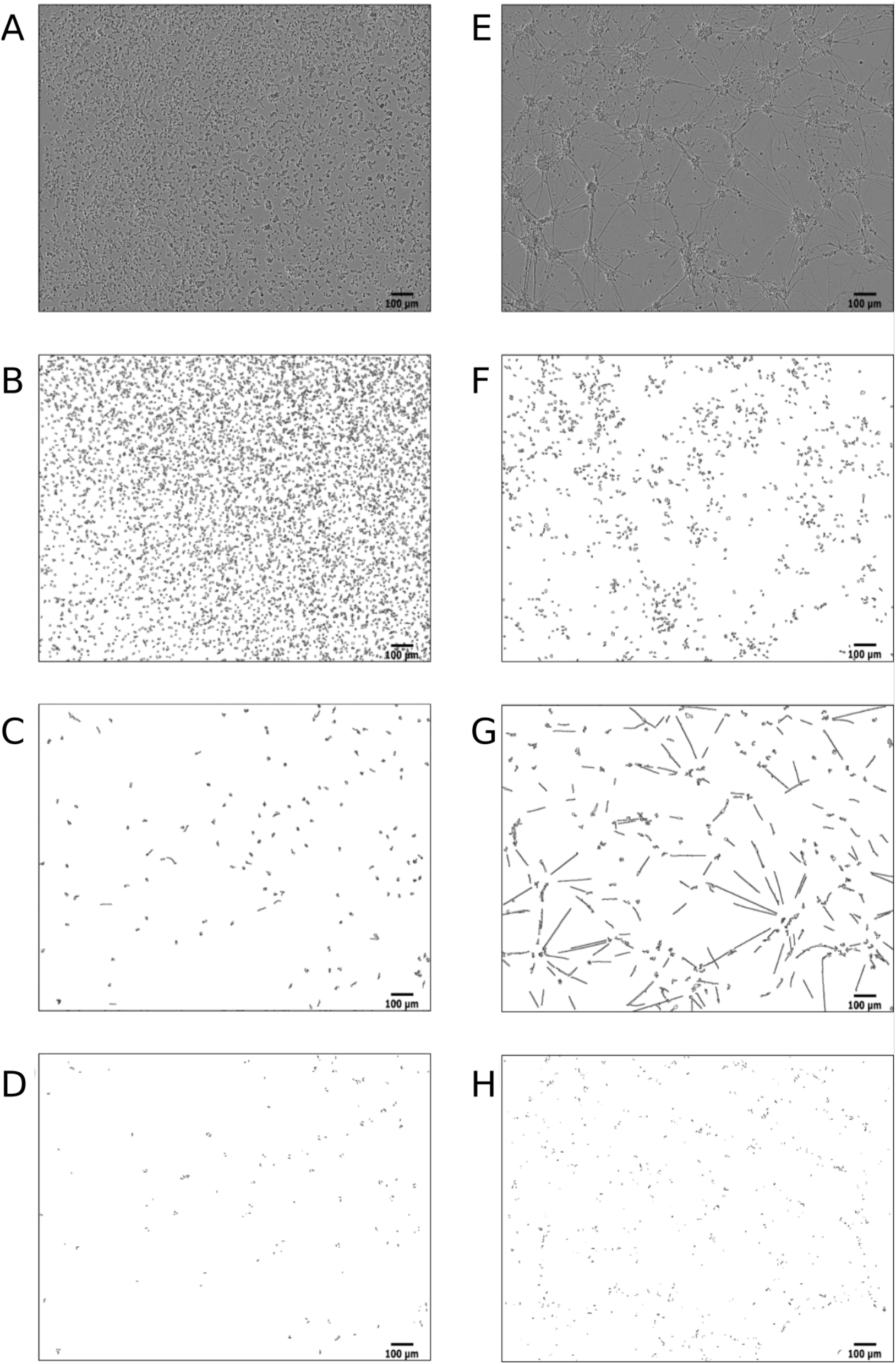
Identified cell structures from ANDA image analysis of CGN cells. (A) Phase contrast image of freshly plated cells. (B) Outlines of identified cell bodies in freshly plated cells. (C) Outlines of identified neurites in freshly plated cells. (D) Outlines of identified neurite attachment points in freshly plated cells. (E) Phase contrast of CGN cells at day in vitro (DIV) 3. (F) Outlines of identified cell bodies at DIV 3. (G) Outlines of identified neurites at DIV 3. (H) Outlines of identified neurite attachment points at DIV 3. Scale bar is 100 micrometres.

### Measurement of neuronal morphology in cell models across species

We next used ANDA to measure the morphological dynamics in chicken CGN, rat PC12N and human NT2N cells. All three models exhibit morphologies applicable for quantification of neuronal metrics. CGNs and NT2Ns exhibited a decrease in cell body count throughout differentiation, whereas PC12Ns displayed an initial slight increase up to 50 hours followed by a decrease (Figure 3). The drop in PC12Ns after 50 hours can be explained by the developed dependency to nerve growth factor (NGF), and the subsequent depletion thereof in the culture media. Mean neurite lengths remained stable with a slight increase for all three models (Figure 3A, C and E). Overall, the number of cell bodies decreased in the CGN model, however there may also be some cell bodies that cluster together that is difficult to distinguish with an automated workflow. The NT2N model exhibited decrease in numbers of cell bodies whereas cell bodies for PC12Ns showed an increase and later decreased towards the end of the experiment.

**Figure 3.**
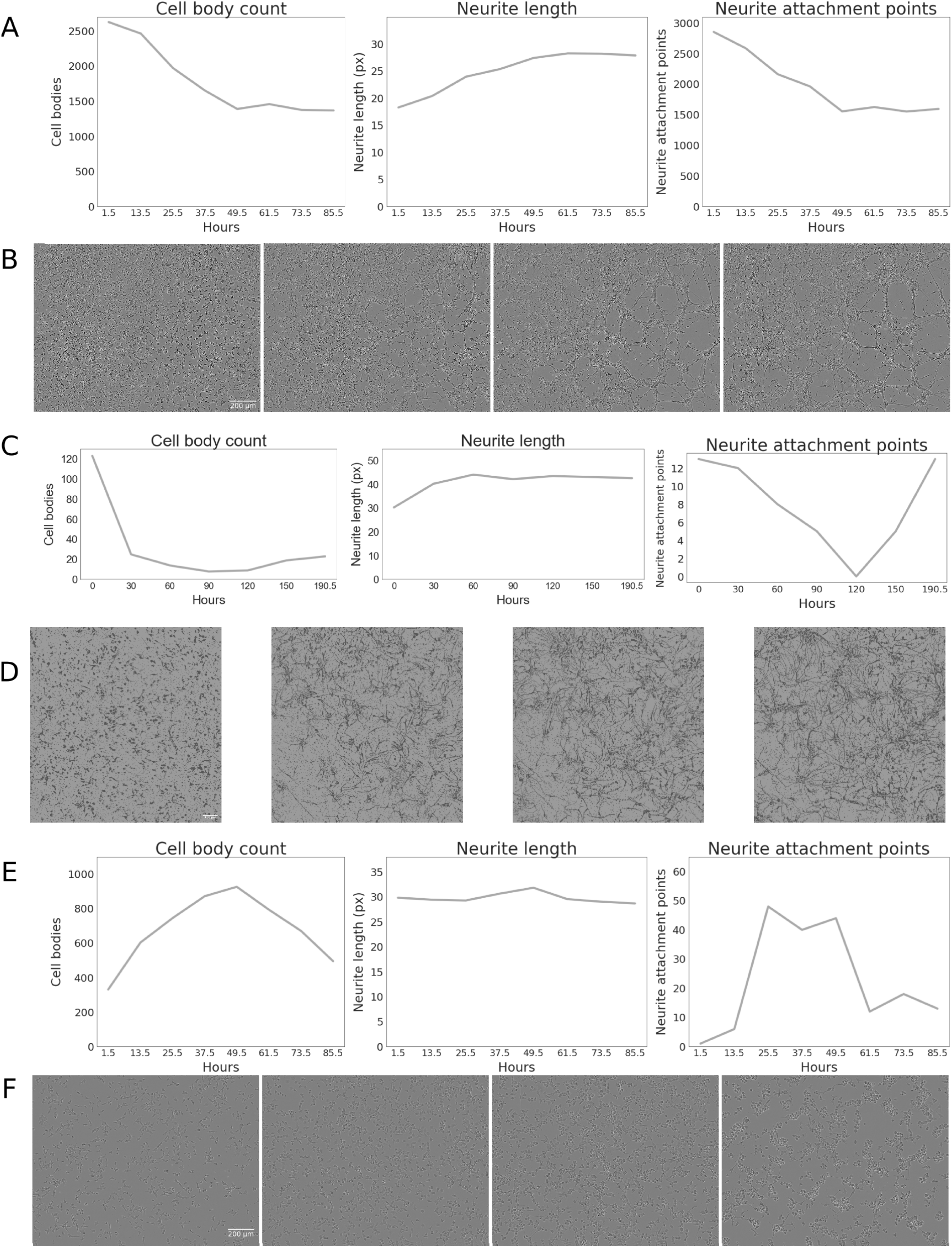
Analysis of neuronal differentiation metrics in CGNs, NT2Ns and PC12Ns. Time course measurements of neuronal differentiation in (A) CGNs, (C) NT2Ns and (E) PC12Ns. Sample phase contrast images of (B) differentiating CGNs (at 1.5, 25.5, 49.5 and 85.5 hours) (D) Weka segmented images of differentiating NT2N cells (at 0, 60, 120 and 190.5 hours) and (F) differentiating PC12Ns (at 1.5, 25.5, 49.5 and 85.5 hours). Scale bar is 200 µm.

We observe that CGN cells developed longer and fewer neurites over time whereas the NT2N cells extended more and longer neurites (Figure 3A-D). For the PC12N cells we did not detect a substantial difference, which may be because naïve PC12 cells have neurite-like filopodial protuberances that can be difficult to differentiate from neurites using our automated workflow (Figure 3E-F). As fewer cells and neurites indicate fewer cell-to-neurite connections, the frequency of neurite attachment points also decreased in the CGNs. The number of neurite attachment points was slowly decreasing and stabilizing after approximately 50 hours in the CGN cells (Figure 3A). Similarly, the number of neurite attachment points slowly decreased but after approximately 120 hours followed by a steady increase in the differentiating NT2N cells (Figure 3C). The number of neurite attachment points curve for PC12N cells had a rapid increase followed by stabilization and decrease (Figure 3E). Both PC12N and NT2N cells presented a low number of neurite attachment points compared to CGNs.

### Quality assessment by manual quantification

To verify the image analysis quality of ANDA, we have used a selection of images of CGN at early and late stages after seeding in vitro and compared the quantified output from ANDA with manual measurements (Figure 4). The comparison between manual quantification and ANDA’s ability to quantify cell bodies indicates that ANDA tends to underestimate the true number of cell bodies in each image. However, both analyses showed the same trend of decreasing number of cell bodies over time (Figure 4A). We next compared the neurite length analysis output from ANDA with manual measurements and the commercially available IncuCyte® NeuroTrack Software Module (NeuroTrack) provided by Essen BioScience (Sartorius). All three analysis methods showed a similar trend in increasing mean neurite lengths over time, but both computational methods tended to underestimate the neurite lengths, especially at the later time points of CGN compared to manual counts (Figure 4B).

**Figure 4.**
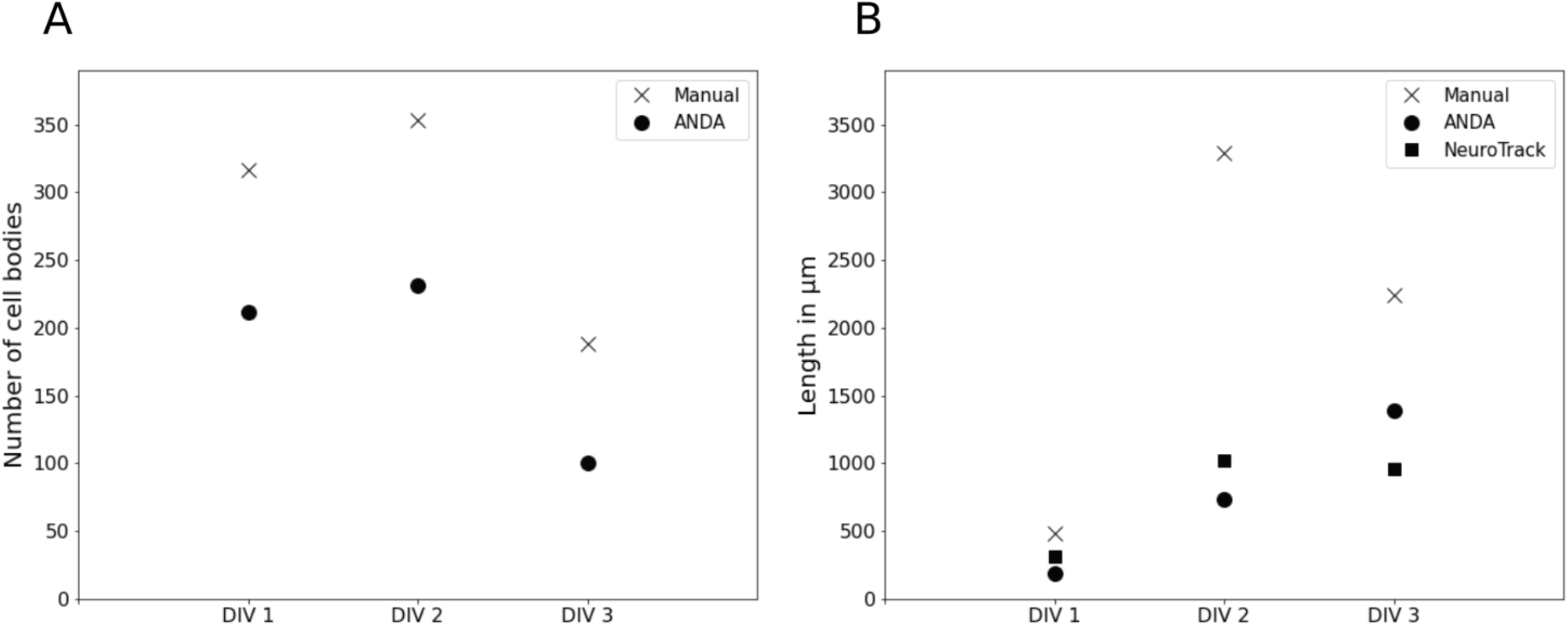
Comparison of analysis with ANDA to NeuroTrack and manual quantification of CGN cells. (A) Comparison of output after analysis of cell body count with ANDA and manual quantification. (B) Comparison of output after analysis of neurite length with ANDA, manual measurements and NeuroTrack. Manual measurements are presented as an average of n = 2 independent measurements at three stages of in vitro differentiation (DIV).

Comparison of cell body analytics between ANDA and NeuroTrack could not be performed quantitatively, as NeuroTrack quantifies cells into clusters instead of attempting to delineate individual cells (Figure 5). Judging qualitatively, both ANDA and NeuroTrack showed a tendency to detect cell bodies with minor differences to the manual measurements in the early and mid-stage time points (Figure 5A and B). At the latest time point, NeuroTrack identified the density of cell bodies more precisely than ANDA (Figure 5C), while yielding more false positives. The number of neurite structures was underestimated by both computational methods especially at the mid-stage time point (Figure 5B). Although neurite counts and lengths were underestimated in comparison to the manual quantifications thereof, both ANDA and NeuroTrack tended to correctly identify oblong structures as neurites during the latest time point in CGN development (Figure 5C).

**Figure 5.**
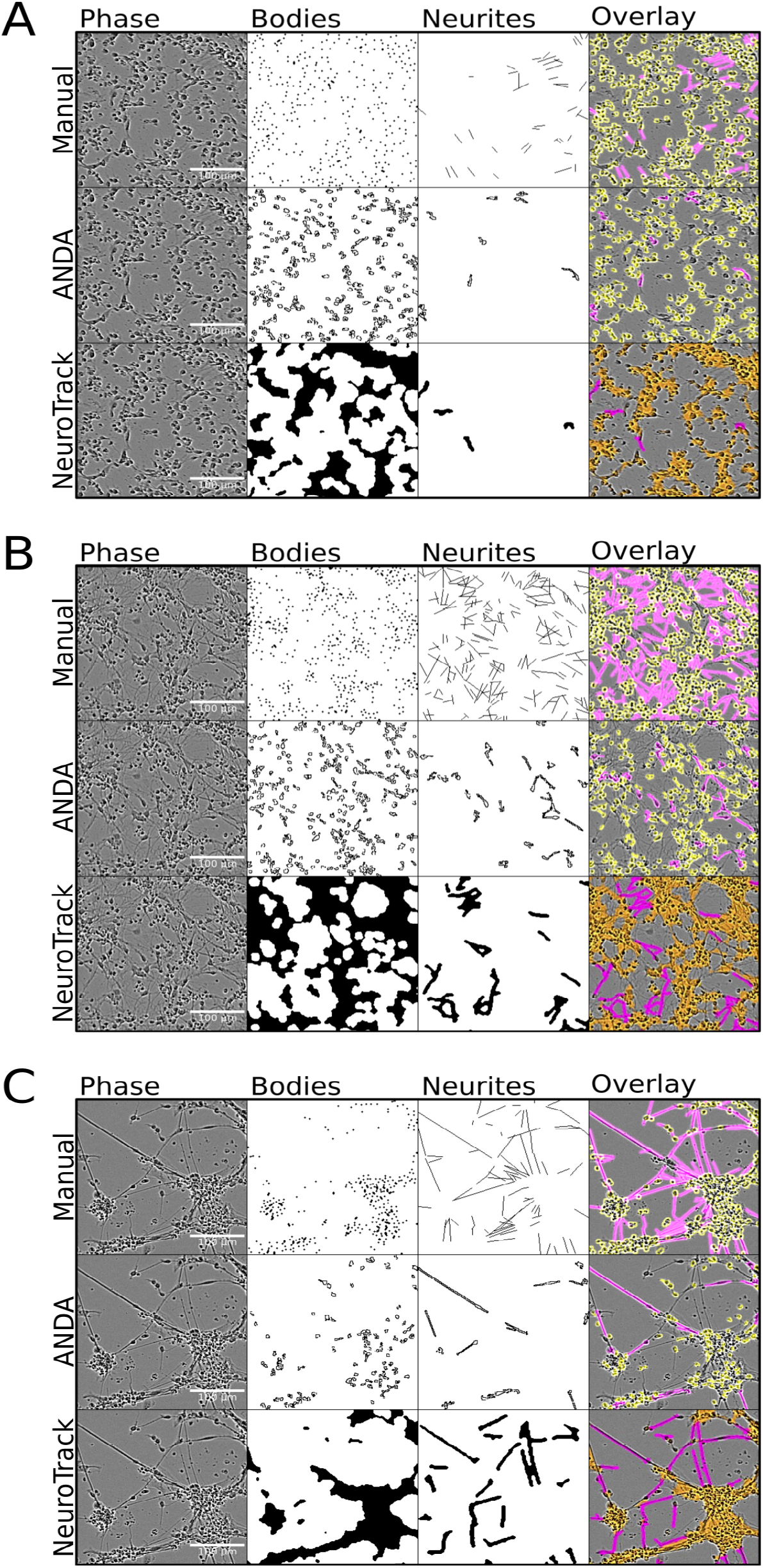
Mask comparisons between manual measurements, ANDA and NeuroTrack. From left to right: Phase contrast images, cell bodies, neurites, mask overlay. From top to bottom: manual measurements, ANDA, NeuroTrack. (A) CGNs day in vitro (DIV) 1; (B) DIV 2 and (C) DIV 3.

## Discussion

ANDA is an image analysis tool particularly useful for analysis of large datasets from live cell high-throughput neuronal image time-series. The comparison of the metrics of the cell models suggests that the numerical output from analysis with ANDA is relatively consistent with what is observed in the images. In comparing ANDA to NeuroTrack, the two methods yielded relatively similar outputs. However, whereas NeuroTrack averages cell body counts into clusters, ANDA quantifies individual soma, including somatic characteristics such as size and shape, yielding a much higher granularity of data. The discrepancies between the numerical output and what is observed in the images can to a large degree be attributed to identification of false positive structures such as dead cell debris or structures falsely quantified due to oversegmentation of the images. As with any automated method, ANDA comes with some limitations.

These limitations are mostly dependent on the quality of the data, with higher quality reducing the possibility of errors by image pre-processing and segmentation. Therefore, we have outlined some recommendations for image analysis to improve the quality of the output. As ANDA is not performing uncertainty assessment, we recommend the use of images with notable contrast between cells and background. If low-contrast cells are used, we strongly recommend Weka-based pre-segmentation prior to analysis with ANDA[32]. The experimental setup should ensure that cell densities do not become excessively high. At too high cell densities ANDA is not able to correctly segment cells from each other or the background. Furthermore, ANDA is not able to distinguish individual neurites in fasciculated neurite bundles and will only retrieve the length of the identified oblong objects, regardless of its width. False positive neurites can be identified by comparing image outlines with the raw data. ANDA also includes the option to set an aspect ratio threshold for which oblong structures are regarded as false positive or not. Primary cells, glial cells and neuronal cells cannot be clearly distinguished by phase contrast alone. The user can, however, isolate granule cells from the images based on criteria such as size and shape.

We have shown that ANDA was able to capture the overall trend from the images represented here, albeit with a subtle over- or under-estimation of the number of neuronal metrics (Figure 3). To address this, we have included an option to remove falsely identified neurites from the final output based on neurite aspect ratio in ANDA. This can be achieved by setting a certain threshold. All neurite structures with an aspect ratio below this will be regarded as false positive and therefore not included in the final output when mean neurite length and number of neurite attachment points are summarized.

## Conclusion

We have demonstrated that the open-source tool ANDA is suitable for analysis of high-throughput images of differentiating neuronal cells from mouse, rat, and chicken in vitro models. ANDA can effectively analyse time-series image sets of differentiating neuronal cells with vastly differing morphologies. To that end, we have shown that ANDA is an accurate, versatile, efficient, and user-friendly tool for quantification of neuronal morphometrics in different model systems.

**Project name:** ANDA: An open-source tool for automated image analysis of neuronal differentiation

**Project home page:** https://github.com/EskelandLab/ANDA and https://www.nitrc.org/projects/anda_neuronal/

**Operating system(s):** Linux, MacOS, Windows

**Programming language:** Python, shell, HTML, CSS, JavaScript, Rust

**License:** MIT

**Any restrictions to use by non-academics:** MIT

## Availability of data and materials

Example datasets generated and analysed during the current study are available as downloads in the NeuroImaging Tools & Resources Collaboratory (NITRC) https://www.nitrc.org/projects/anda_neuronal.

## Acknowledgements

We thank Ajay Yadav, Section for Pharmacology and Pharmaceutical Biosciences for the PC12N images and the National Institute of Occupational Health in Norway for sharing of IncuCyte® ZOOM equipment. The majority of informatic analysis was performed at saga super computing resources (Project NN9632K) provided by UNINETT Sigma2—the National Infrastructure for High Performance Computing and Data Storage in Norway. We have featured ANDA and deposited microscopy images used in this study in https://www.nitrc.org. NITRC, NITRC-IR, and NITRC-CE have been funded in whole or in part with Federal funds from the the Department of Health and Human Services, National Institute of Biomedical Imaging and Bioengineering, the National Institute of Neurological Disorders and Stroke, under the following NIH grants: 1R43NS074540, 2R44NS074540, and 1U24EB023398 and previously GSA Contract No. GS-00F-0034P, Order Number HHSN268200100090U.

