## Supplemental Data 1 for "ANDA: An open-source tool for automated image analysis of neuronal differentiation"

### ANDA: An open-source tool for neuronal differentiation analysis from high throughput imaging experiments.

### Supplementary Figures

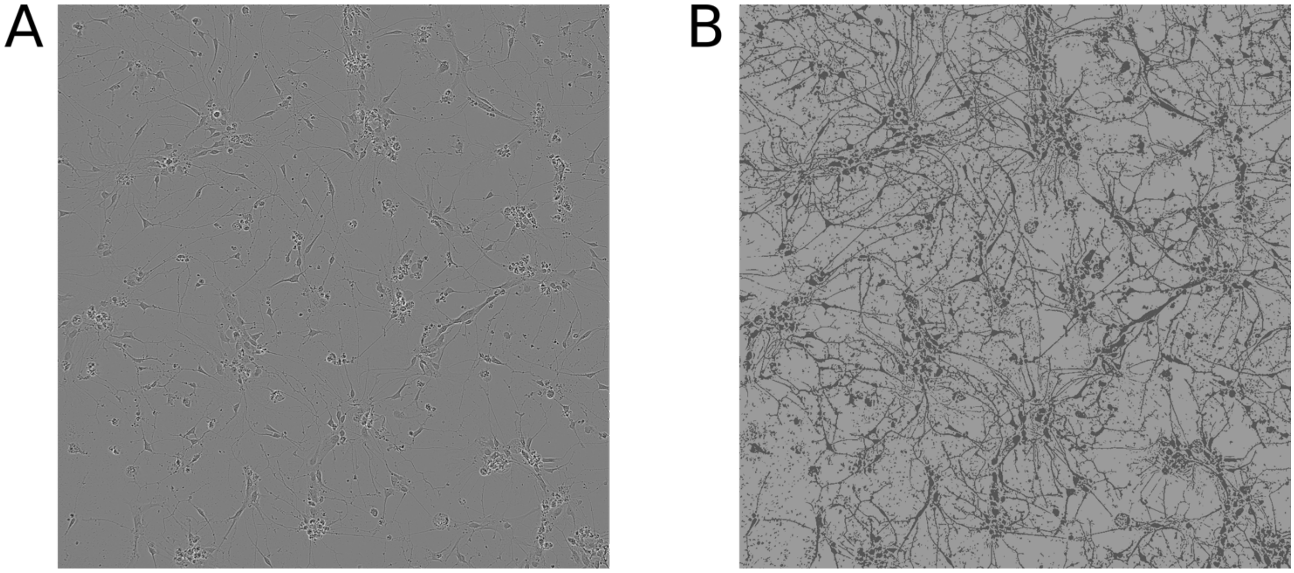


**Supplementary figure 1. NT2N cells before and after segmentation with Weka.** A: Phase contrast image of NT2N obtained from Incucyte® S3. B: The same image after Weka segmentation. Scale bar is 200 μm.

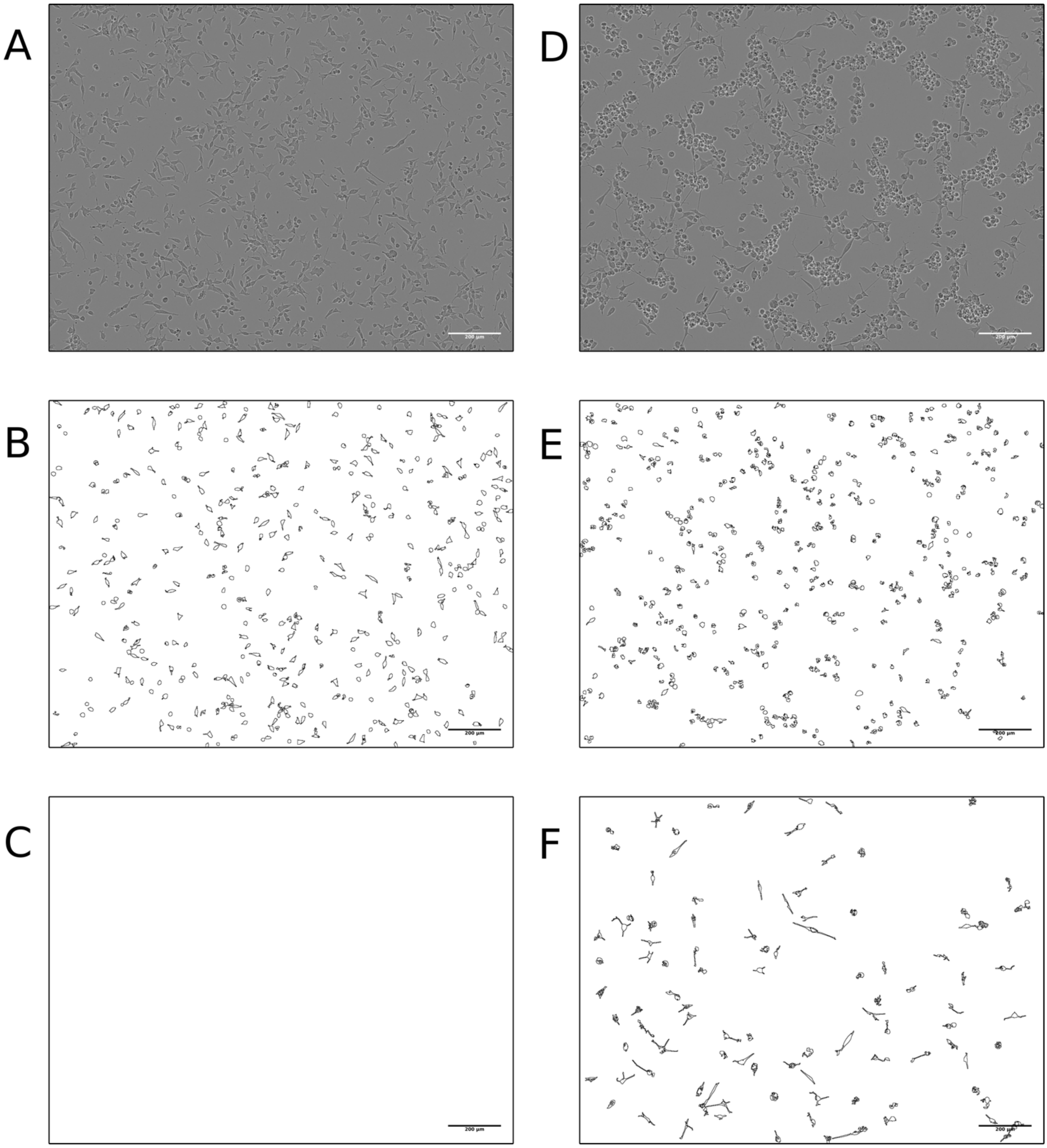

**Supplementary figure 2. Identified cell structures from ANDA image analysis of PC12N cells.** (A) Phase contrast image of freshly plated cells. (B) Outlines of identified cell bodies in freshly plated cells. (C) Outlines of identified neurites in freshly plated cells. (D) Phase contrast of PC12N cells at day 3. (E) Outlines of identified cell bodies at day 3. (F) Outlines of identified neurites at day 3. Scale bar is 200 micrometres.

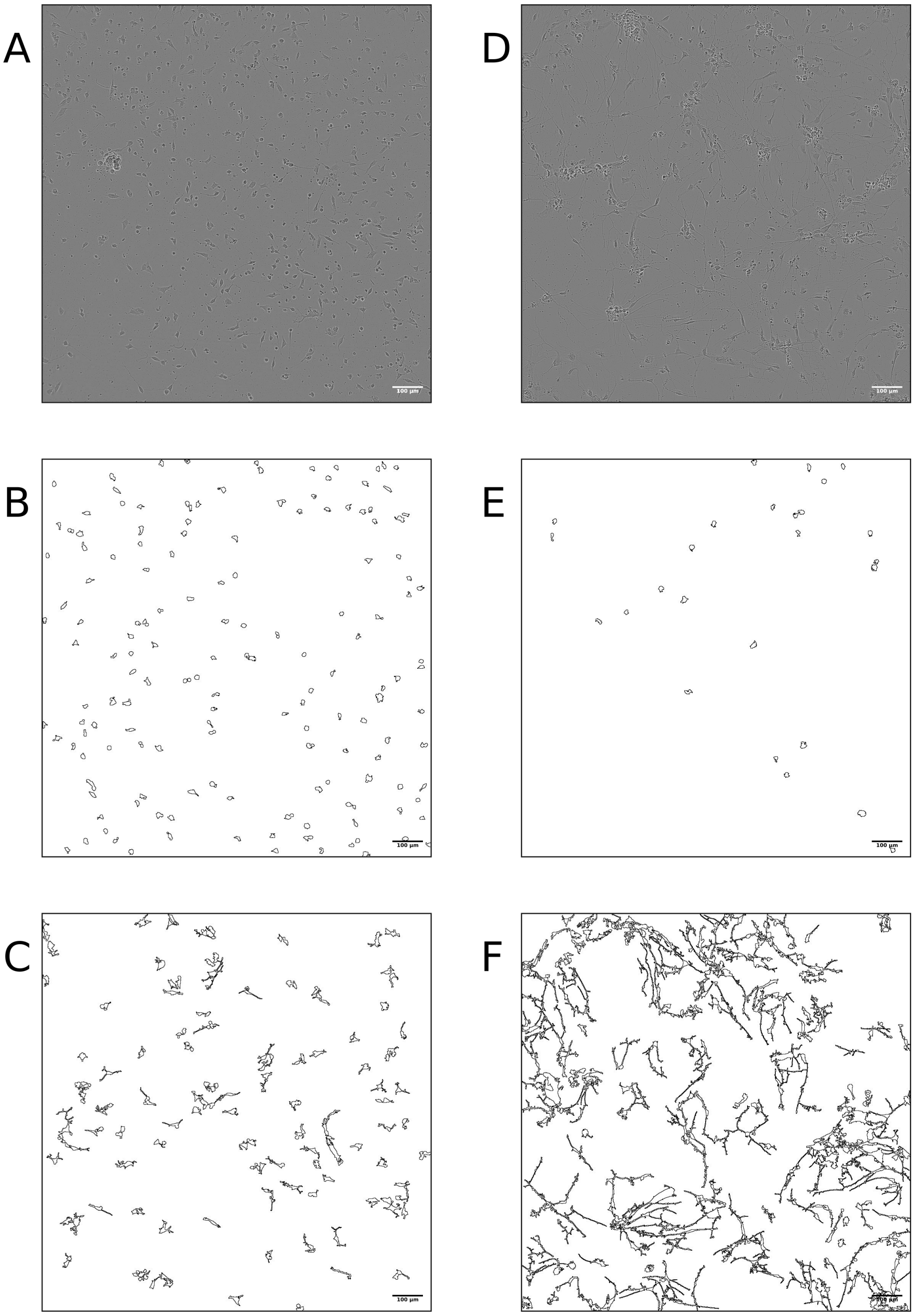


**Supplementary figure 3. Identified cell structures from ANDA image analysis of Weka segmented NT2N cells.** (A) Weka segmented image of freshly plated cells. (B) Outlines of identified cell bodies in freshly plated cells. (C) Outlines of identified neurites in freshly plated cells. (D) Weka segmented image at neuronal differentiation day 3. (E) Outlines of identified cell bodies at day 3. (F) Outlines of identified neurites at day 3. Scale bar is 200 micrometres.

### Supplementary Table

**Supplementary Table 1.** Size and shape criteria used for analysis of NT2Ns, CGNs and PC12Ns for ANDA.

| | Min. circularity | Max. circularity | Min. size (px) | Max. size (px) |
|---|---|---|---|---|
| | Cell bodies | | | |
| NT2Ns | 0.38 | 1.00 | 90 | 900 |
| CGNs | 0.40 | 1.00 | 16 | 175 |
| PC12Ns | 0.20 | 1.00 | 60 | 175 |
| | Neurites | | | |
| NT2Ns | 0.00 | 0.38 | 200 | 1800 |
| CGNs | 0.00 | 0.38 | 89 | 1249 |
| PC12Ns | 0.00 | 0.22 | 160 | 1000 |

### Supplementary methods

#### Chicken cerebellar granule neuronal cells

The generation of chicken granule neuronal cells were performed as described previously [1]. Fertilized chicken eggs (*Gallus gallus*) were incubated at 37.5 °C, 45 % humidity for 17 days, anaesthetized and sacrificed prior to cerebellar excision (approved under Norwegian Food Safety Authority FOTS ID 13896). CGN culture was prepared by trypsinization and trituration of pooled cerebella, followed by seeding in Basal Eagle's Medium supplemented with chicken serum into PLL-coated 96-well plates at 530.000 cells/ $cm^2$. Following overnight incubation, serum-containing medium was replaced with defined medium supplemented with 10 µM cytosine β-D-arabinofuranoside to limit glial proliferation.

#### *In vitro* differentiation of NTERA2 and PC12

The generation of neuronal NT2Ns was performed as described previously [1]. Briefly, human NTERA2 embryocarcinoma cells (ATCC, UK) were seeded into 100 mm bacterial dishes at $5\times10^5$ cells/mL in 10 mL Dulbecco's modified Eagle's medium supplemented with 10% foetal bovine serum and incubated at 37°C with 5 % $CO_2$ on a rotator for 2-4 days until spheroids formed. Spheroids were differentiated in rotary culture in serum-containing medium supplemented with 10 µM retinoic acid for 6 days, with half-volume medium change taking place every two days. On day 6, serum-containing medium was replaced with defined medium consisting of DMEM/F12 supplemented with B27 and N2 (according to manufacturer's instructions), as well as 10 µM retinoic acid, in which the spheroids were differentiated in rotary culture for another 14 days, with half-volume medium changes taking place every two days. Spheroids were then trypsinized and seeded into PLL- and geltrex-coated 96-well plates at 50.000 cells/$cm^2$ in 1:1 mixture of conditioned and fresh defined medium supplemented with 10 µM retinoic acid.

The rat pheochromocytoma cell line PC12 was cultivated in DMEM supplemented with 5% horse serum and 10% foetal bovine serum as described previously [2]. Cells were seeded into 96-well plates at a density of 8 x $10^3$ cells/$cm^2$. Following overnight incubation, medium was replaced with fresh DMEM containing 2% horse serum and 50 ng/mL nerve growth factor for three days.

**Imaging with Incucyte®**

Live-cell imaging was performed at 37 °C and 5% $CO_2$ using the Incucyte® ZOOM and Incucyte® S3 platforms (EssenBioScience, Hertfordshire, UK). CGN and PC12N experiments were carried out in TPP® 96-well plates (Sigma-Aldrich #Z707910) in Incucyte® ZOOM, whereas NT2N experiments were undertaken in Corning® black-frame clear-bottom 96-well plates (Corning #3603) in Incucyte® S3. Phase contrast images were acquired at 10x magnification using the Incucyte®'s built-in settings corresponding to each plate type.

**Manual count and quantification with Neurotrack**

To evaluate the performance of ANDA, its outputs were compared to the outputs of the EssenBioscience Incucyte® ZOOM Neurotrack Analysis Software Modul, as well as to manual quantifications. Two randomly selected areas of three sets of images of CGNs from DIV1, DIV2, and DIV3 were analyzed using the three different modalities. Manual quantifications were performed by two expert personnel, where cell soma, neurites, and neurite branch points were manually labelled, quantified, and averaged across the two experimenters' outputs. The Neurotrack quantification module was trained using a set of analysis parameters optimized based on a training set from the CGN cells consisting of three images per time-point at 6, 24, 48, and 72 h post-seeding. Analysis with ANDA was executed using the CGN analysis mode.

**Requirements for ANDA**

To be able to use ANDA, the user needs to have and Python 3 installed [3].

3. Van Rossum G, Drake FL. Python 3 Reference Manual. Scotts Valley, CA: CreateSpace.; 2009.
